# Tubular STAT3 limits renal inflammation in autosomal dominant polycystic kidney disease

**DOI:** 10.1101/2019.12.12.873901

**Authors:** Amandine Viau, Maroua Baziz, Amandine Aka, Clément Nguyen, E. Wolfgang Kuehn, Fabiola Terzi, Frank Bienaimé

**Author notes:** Corresponding author: Frank Bienaimé INSERM U1151 Team: Mechanisms and Therapeutic Strategies of Chronic Kidney Disease. Hôpital Necker Enfants Malades Tour Lavoisier, 6ème étage 149 Rue de Sèvres 75015 Paris France.

## Abstract

The inactivation of the ciliary proteins polycystin 1 or 2 leads to autosomal dominant polycystic kidney disease (ADPKD), the leading genetic cause of chronic kidney disease. Both cilia signaling and interstitial inflammation play a critical role in the disease. Yet, the reciprocal interactions between immune and tubular cells are not well characterized. The transcription factor STAT3, which is suspected to fuel ADPKD progression, is involved in crosstalks between immune and non-immune cells in various tissues and is a component of the cilia proteome. Here, we explore how STAT3 intersects with cilia signaling, renal inflammation and cyst growth using conditional murine models of post-developmental *Pkd1*, *Stat3* and cilia ablation. Our results indicate that, although primary cilia directly modulate STAT3 activation *in vitro*, the bulk of STAT3 activation in polycystic kidneys occurs through an indirect mechanism in which primary cilia trigger macrophage recruitment to the kidney, which in turn promotes STAT3 activation. Surprisingly, while disrupting *Stat3* in *Pkd1* deficient tubules slightly reduced cyst burden, it resulted in a massive infiltration of the cystic kidneys by macrophages and T cells, precluding any improvement of kidney function. Mechanistically, STAT3 represses the expression of the inflammatory chemokines CCL5 and CXCL10 in polycystic kidneys and cultured tubular cells. These results demonstrate that STAT3 is not a critical driver of cyst growth in ADPKD but plays a major role in the crosstalk between immune and tubular cells that shapes disease expression.

## INTRODUCTION

Autosomal dominant polycystic kidney disease (ADPKD), the leading genetic cause of kidney disease, accounts for 5% of patients with end-stage renal disease (ESRD). Therapies allowing patients to avoid ESRD are currently lacking. Indeed, the only drug to have shown some efficacy to date only modestly reduces kidney function decline at the expense of considerable side effects^1^. Thus, a better understanding of ADPKD pathophysiology is of major importance to delineate more effective therapies.

ADPKD is caused by inactivating mutations affecting either *PKD1*, which encodes polycystin 1, a putative orphan receptor, or *PKD2*, which encodes polycystin 2, a cation channel that interacts with polycystin 1^2^. Although ADPKD patients carry heterozygous germline *PKD1* (or *PKD2*) inactivating mutations, cyst development is caused by somatic mutations affecting the remaining functional *PKD1* (or *PKD2*) allele in a subset of tubular cells that will proliferate to form cysts^3^. In adult mice, bi-allelic loss of *Pkd1* in tubular cells results in a biphasic proliferative response leading first to non-cystic kidney enlargement (pre-cystic phase), which is followed by the rapid development of cysts (cystic phase)^4,5^. While cyst formation is the consequence of a genetic defect in tubular cells, this process is not cell-autonomous. Indeed, cystogenesis is associated with important modifications of the renal microenvironment, most importantly immune cell recruitment in the vicinity of *Pkd1* deficient tubules, which, in turn, affects cyst growth and kidney function decline^6–10^, as well as quantitative and qualitative alterations of the extracellular matrix^11,12^, which are also thought to contribute to disease progression.

Primary cilia play an essential role in ADPKD^4,10^. The primary cilium is a microtubule-based organelle that protrudes from the apical surface of tubular cells to integrate mechanical and chemical cues delivered by the urinary flow. The Polycystin 1/2 complex localizes to the primary cilium and this localization is essential to repress cyst formation^13,14^. Remarkably, cilia ablation drastically suppressed both pre-cystic cell proliferation and cyst formation in *Pkd1* mutant animals^4,10^. However, the nature of the deleterious signals delivered by cilia in this context are only starting to emerge. We and others have recently shown that cilia ablation prevents the induction of the macrophage chemoattractant CCL2 in response to *Pkd1* disruption and that *Ccl2* inactivation dampens renal macrophages infiltration and cyst formation in *Pkd1* mutant mice^7,10^. While these results demonstrate that primary cilia control the communication between immune and tubular cells, the molecular network shaping the interaction between polycystin deficient tubular cells and the neighboring immune cells remain poorly understood. STAT3 is a pleiotropic transcription factor involved in a large spectrum of physiological and pathophysiologic processes^15^. STAT3 activation is principally mediated by its phosphorylation on tyrosine 705 by various cytosolic or membrane kinases, which promotes its nuclear accumulation and transcriptional activity. STAT3 represents an interesting candidate for immune-tubular cell communication as it stands at the crossroads of cilia signaling, immune regulation and cyst growth. Numerous studies have pinpointed STAT3 as a general driver of both pro- and anti-inflammatory crosstalk between epithelial and neighboring immune cells^16,17^. A recent proximity biotinylation based proteomic screen identified STAT3 as a *bona fide* component of the ciliary machinery^18^. In ADPKD and murine PKD models, phosphorylated STAT3 accumulates in the nucleus of cyst lining epithelial cells^19,20^. Furthermore, the cytosolic tail of polycystin 1 binds STAT3 *in vitro* and modulates its transcriptional activity^18^. Lastly, the positive effect of STAT3 inhibitory drugs in orthologous models of the disease has led to the assumption that tubular STAT3 promoted cystogenesis^19,20^. However, how STAT3 intersects with cilia signaling, renal inflammation and cyst growth in the disease is not understood. The limited specificity of the inhibitors used in these studies^20,21^ and their impact on STAT3 activation in non-tubular cells preclude definitive conclusions regarding the precise function of tubular STAT3 in polycystin deficient tubular cells.

Considering these uncertainties, we aimed to clarify (i) the function of primary cilia in STAT3 activation in tubular cells and (ii) its subsequent function in cyst growth, renal inflammation and kidney function decline.

## MATERIALS AND METHODS

### Mice

All animal experiments were conducted according to the guidelines of the National Institutes of Health Guide for the Care and Use of Laboratory Animals, as well as the German and French laws for the animals welfare, and were approved by regional authorities. Mice were housed in a specific pathogen-free facility, fed ad libitum and housed at constant ambient temperature in a 12-hour day/night cycle. Breeding and genotyping were done according to standard procedures.

*Pkd1*^flox/flox^ mice (B6.129S4-Pkd1tm2Ggg/J, stock number: 010671, C57BL/6 genetic background) and *Ccl2-RFP*^flox/flox^ (B6.Cg-Ccl2tm1.1Pame/J, stock number: 016849, C57BL/6 genetic background) were purchased from The Jackson Laboratories, *Kif3a*^flox/flox^ mice (Kif3atm1Gsn, C57BL/6 genetic background) were kindly provided by Peter Igarashi, and were crossed to Pax8rtTA (Tg(Pax8-rtTA2S*M2)1Koes)^22^ and TetOCre (Tg(tetO-cre)1Jaw)^23^ mice to generate an inducible tubule-specific *Pkd1* knockout (further referred to as i*Pkd1*^ΔTub^), *Pkd1*; *Ccl2* knockout (further referred to as i*Pkd1*^ΔTub^; i*Ccl2*^ΔTub^) and *Pkd1*; *Kif3a* knockout (further referred to as i*Pkd1*^ΔTub^; i*Kif3a*^ΔTub^) as previously described^10^. *Stat3*^flox/flox^ mice (Stat3tm1Vpo1, initially kindly provided by Valeria Poli, FVB/N genetic background)^24^ were backcrossed for three generations with i*Pkd1*^ΔTub^ mice on a pure C57BL/6 genetic background. The progeny was then intercrossed to generate mice with inducible tubule-specific *Pkd1* knockout (further referred as i*Pkd1*^ΔTub^) or *Pkd1; Stat3* knockout (further referred to as i*Pkd1*^ΔTub^; i*Stat3*^ΔTub^) on a common mixed FVB/N; C57BL/6 genetic background.

To induce floxed alleles recombination mice received doxycycline (Abcam, ab141091) in drinking water (2mg/mL with 5% sucrose, protected from light) from post-natal day 28 (P28) to P42. Littermates (lacking either TetOCre or Pax8rtTA) were used as controls. Experiments were conducted on males.

### Plasma Analyses

Retro-orbital blood was collected from anaesthetized mice. Plasma blood urea nitrogen (BUN) and plasma creatinine was measured using a Konelab 20i analyzer (Thermo Scientific).

### Morphological Analysis

Mouse kidneys were fixed in 4% paraformaldehyde, embedded in paraffin, and 4µm sections were stained with periodic acid-Schiff (PAS). Stained sections were imaged using an E800 microscope (Nikon) equipped with a Plan Neofluar 20x/0.30 NA objective and a digital Camera Dx/m/1200 (Nikon) coupled to NIS software (Laboratory Imaging Ltd). PAS stained full size kidney images were recorded using a whole slide scanner Nanozoomer 2.0 (Hamamatsu) equipped with a 20x/0.75 NA objective coupled to NDPview software (Hamamatsu).

Cortical tubular lumen fractional area was measured with ImageJ software from PAS stained full size kidney images. Interstitial infiltration score was evaluated by two independent observers in a blinded fashion assessing the overall infiltration of the whole kidney section stained with PAS.

### Cell culture

Madin-Darby canine kidney (MDCK, kind gift of Prof. Kai Simons, MPI-CBG, Dresden, Germany) cells were cultured using Dulbecco’s Modified Eagle Medium supplemented with 10% fetal bovine serum (Sigma-Aldrich, F7524) and 1% Penicillin-Streptomycin (Life Technologies, 15140122). MDCK cells with a tetracycline-inducible *Kif3a* knockdown (further referred to as Kif3a-i), *Ift88* knockdown (further referred to as Ift88-i) or control (further referred to as Luci-i) using a lentivirus-based transduction system have been previously described^10^. 150,000 cells/cm² were seeded with or without 1µg/mL doxycline hyclate (Abcam, ab141091) for 10 days on 12mm Transwell polycarbonate membranes (COSTAR, 3401). RNAs and proteins were extracted after 16 hours serum deprivation.

Mouse inner medullary collecting duct (mIMCD-3) cells were grown in 50% DMEM high-glucose pyruvate (Gibco, 41966052), 50% F-12 Nutrient Mixture (Gibco, 21765037) supplemented with 10% fetal bovine serum and 1% Penicillin-Streptomycin. mIMCD-3 cells with a stable *Stat3* knockdown (further referred to as Stat3 sh1 and 2) have been already reported^24^. 30,000 cells/cm² mIMCD-3 were seeded. Confluent mIMCD-3 cells were serum deprived for 16 hours and then stimulated with 1µg/mL LPS (Invivogen, tlrl-3pelps). RNAs and proteins were extracted 24 hours after stimulation.

All cells were regularly tested for mycoplasma contamination and were mycoplasma free.

### Quantitative real-time PCR (qRT-PCR)

Total RNAs were obtained from whole kidneys or cells using NucleoSpin^®^ RNA Kit (Macherey Nagel) and reverse transcribed using High-Capacity cDNA Reverse Transcription kit (ThermoFisher Scientific). Quantitative PCR was performed with iTaq™ Universal SYBR® Green Supermix (Bio-Rad) on ViiA 7 Real-Time PCR system (ThermoFisher Scientific) coupled to QuantStudio™ software V1.3. Each biological replicate was measured in technical duplicates. The primers used for qRT-PCR are listed in Supplementary Table 1.

### Immunohistochemistry

4µm sections of paraffin-embedded kidneys were submitted to antigen retrieval and avidin/biotin blocking (Vector, SP-2001). Sections were incubated with primary antibody followed by biotinylated antibody, HRP-labeled streptavidin (Southern Biotech, 7100-05, 1:500) and 3-3′-diamino-benzidine-tetrahydrochloride (DAB, Dako, K3468) revelation. Stained sections were imaged using an E800 microscope (Nikon) equipped with a Plan Neofluar 20x/0.30 NA objective and a digital Camera Dx/m/1200 (Nikon) coupled to NIS software (Laboratory Imaging Ltd).

### Western blot

Kidneys were lyzed in 50mM Tris pH8, 200mM NaCl, 1mM EDTA, 1mM EGTA, 1mM DTT, 1% SDS buffer. Cells were lysed in modified RIPA buffer lysis buffer (150 mM NaCl, 50 mM Tris-HCl pH 7.5, 1% Triton X-100, 0.5% sodium deoxycholate, 0.1%sodium laury sulfate. Nucleo-cytoplasmic fractionation was performed using NePer kit (Thermofisher). Lysis buffers were supplemented with protease and protein phosphatase inhibitors tablets (Pierce, A32959). Protein content was determined with Pierce BCA protein assay kit (Pierce, A23225). Equal amounts of protein were resolved on 4-20% gradient gels (Bio-Rad) under reducing conditions, transferred and incubated with primary and secondary antibodies and visualized using ChemiDoc™ (Bio-Rad) ismaging system coupled to Image Lab software (Bio-Rad).

### Antibodies

#### For western blotting

pSTAT3^Y705^ (Cell Signaling Technology, 9145, 1:1,000), STAT3 (Cell Signaling Technology, 9132, 1:1,000), GAPDH (Millipore, MAB374, 1:5,000), α-TUBULIN (Sigma, T5168, 1:20,000), Lamin A/C (Abcam, ab108922, 1:1,000) and RELA (Cell Signaling Technology, 4764, 1:1,000).

#### For immunostaining

pSTAT3^Y705^ (Cell Signaling Technology, 9145, 1:100), STAT3α (clone D1A5, Cell Signaling Technology, 8768, 1:100), F4/80 (Clone Cl:A3-1, Bio-Rad, MCA497R, 1:100), CD3 (Abcam, ab16669; 1:100) and LY6B (Abcam, ab53457, 1:100).

### Statistical Analysis

Data were expressed as means or as means ± SEM. Differences between the experimental groups were evaluated using one or two-way ANOVA as appropriate, followed when significant (*P* < 0.05) by the Tukey-Kramer test. When only two groups were compared, two-tailed *t* test or Mann–Whitney test was used as appropriate. For Kaplan Meier survival curves, Log-rang test was applied. The statistical analysis was performed using GraphPad Prism V6 software. No samples were excluded from analyses. All image analyses (immunohistochemistry, immunofluorescence) and mouse phenotypic analyses were performed in a blinded fashion.

## RESULTS

### Primary cilia regulate STAT3 activation in tubular cells

To investigate if STAT3 activation occurred at an early stage of the disease, we took advantage of transgenic mice allowing the inducible inactivation of *Pkd1* in tubular cells upon doxycycline treatment (i*Pkd1*^Δtub^). At twelve week of age (6 weeks after the completion of doxycycline treatment), tubular cell proliferation is increased and kidneys are enlarged but minimally cystic^10,25^. Immunolabelling of kidney sections from 12 week old i*Pkd1*^Δtub^ and control mice with STAT3 phospho-specific antibody directed against Tyrosine 705 revealed an increase in both tubular and interstitial phosphorylated STAT3 positive nuclei in polycystin deficient kidneys compared to controls (Figure 1A-C). Of note, most of the signal was observed in areas with mild tubular dilation and interstitial infiltration. Ablation of primary cilia in i*Pkd1*^Δtub^ mice through simultaneous deletion of *Kif3a* markedly blunted both tubular and interstitial STAT3 activation (Figure 1A-C).

**Figure 1:**
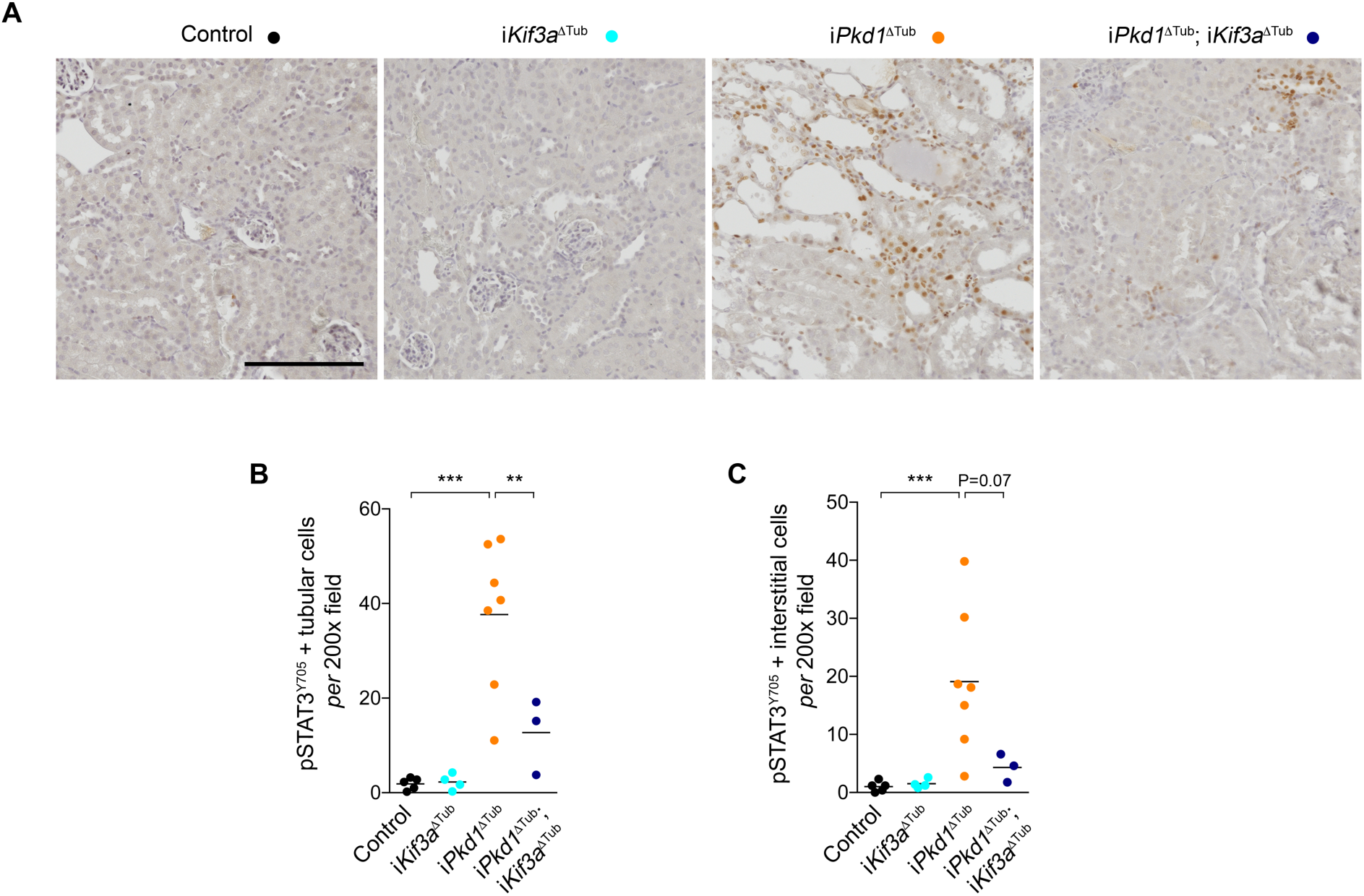
Cilia ablation reduces STAT3 activation in response to *Pkd1* inactivation. (A-C) Representative images **(A)** and quantification **(B-C)** of pSTAT3^Y705^ immunostaining in kidneys from 12 weeks old control, i*Kif3a*^ΔTub^, i*Pkd1*^ΔTub^, and i*Pkd1*^ΔTub^; i*Kif3a*^ΔTub^ mice (6 weeks after the completion of doxycycline treatment). Scale bar: 50µm. Each dot represents one individual mouse. One-way ANOVA followed by Tukey-Kramer test, \*\**P*<0.01, \*\*\**P*<0.001.

We then investigated the mechanisms allowing cilia dependent STAT3 activation *in vivo*. While the identification of STAT3 as a component of the primary cilia proteome renders a direct regulation of STAT3 by primary cilia plausible, primary cilia have been shown to orchestrate an inflammatory response through the induction of CCL2 in tubular cells. In the heart, CCL2 induces macrophage recruitment, which in turn activates STAT3 in cardiomyocytes through paracrine signaling^26,27^. We evaluated the possibility of such a non-cell-autonomous pathway to tubular STAT3 activation *in vivo*. We analyzed the impact of tubule specific *Ccl2* disruption on STAT3 in the polycystic kidney. We and others have previously shown that *Ccl2* disruption reduced renal macrophages infiltrates and cyst burden in polycystic kidneys^7,10^. Indeed, western blot of kidney homogenates revealed that *Ccl2* inactivation strongly reduced renal STAT3^Y705^ phosphorylation (Figure 2A). In line with this observation, *Ccl2* inactivation also reduced the mRNA abundance of the STAT3 transcriptional targets *Socs3* and *Lcn2* (Figure 2B, C). Immunolabelling confirmed that *Ccl2* inactivation markedly reduced both interstitial and tubular STAT3 phosphorylation (Figure 2D-F). However, residual STAT3 activation in renal tubules was detected in the kidneys of i*Pkd1*^ΔTub^; i*Ccl2*^ΔTub^ mice, although to a level that was not statistically different from control kidneys. To investigate if primary cilia can directly modulate STAT3 activation independently of macrophage recruitment, we analyzed STAT3 phosphorylation in tetracycline-inducible *Kif3a*-silenced (Kif3a-i) or *Ift88*-silenced (Ift88-i) MDCK cells. We found that tetracycline treatment, which results in cilia ablation in both cell lines^28,29^, modestly but significantly reduced STAT3 phosphorylation as compared to control cells expressing an inducible shRNA targeting luciferase (Luci-i; **Supplementary Figure 1A, B**).

**Figure 2:**
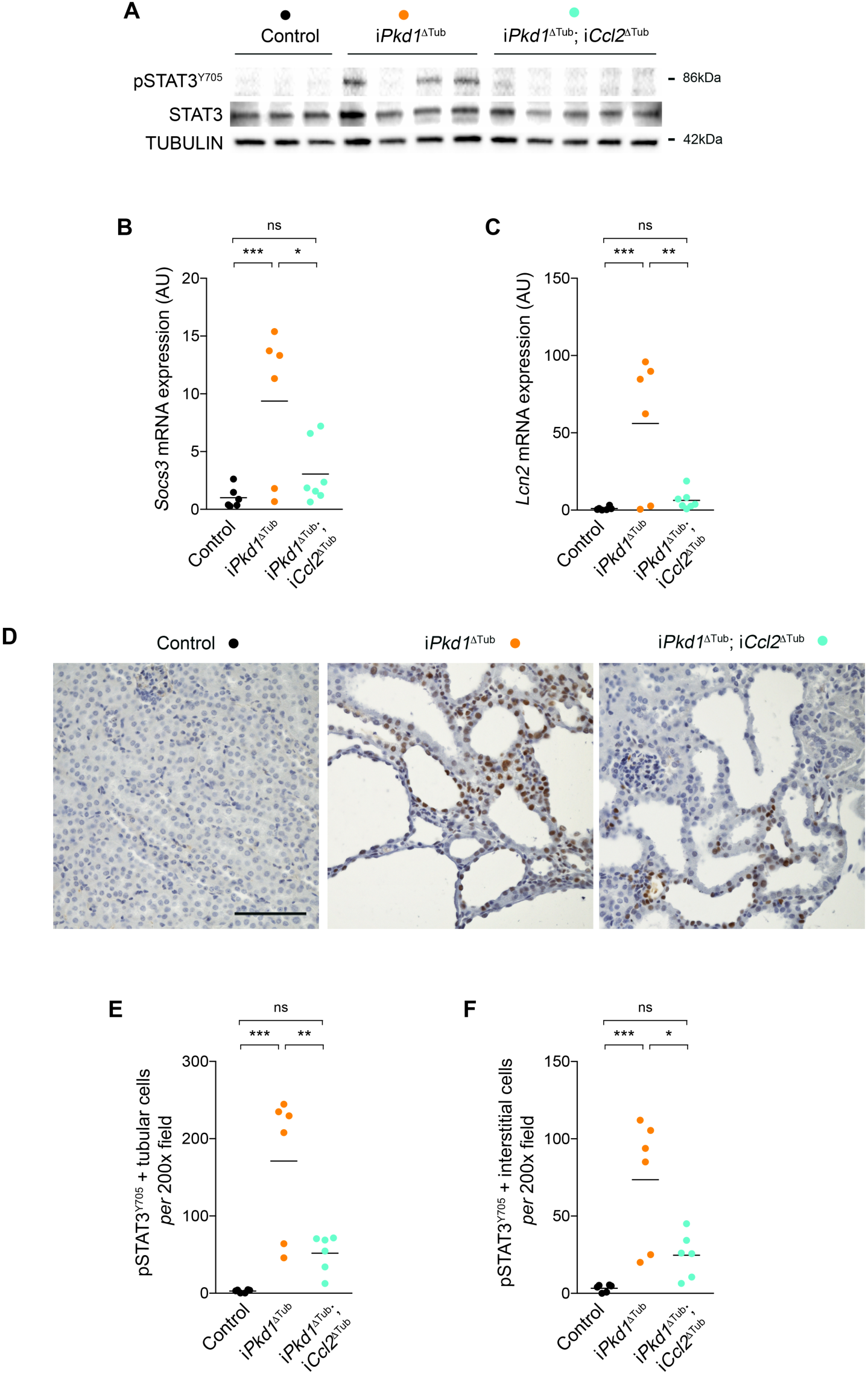
*Ccl2* expression by tubular cells is required for STAT3 activation in response to *Pkd1* disruption. **(A)** Western blot of STAT3 phosphorylation in kidney lysates from 13 weeks old control, i*Pkd1*^ΔTub^ and i*Pkd1*^ΔTub^; i*Ccl2*^ΔTub^ mice (7 weeks after the completion of doxycycline treatment). Each lane indicates one individual mouse. **(B-C)** Quantification of *Socs3* **(B)** and *Lcn2* **(C)** mRNA abundance in kidneys from 13 weeks old control, i*Pkd1*^ΔTub^ and i*Pkd1*^ΔTub^; i*Ccl2*^ΔTub^ mice. AU: arbitrary units. **(D-F)** Representative images (**D**) and quantification **(E-F)** of pSTAT3^Y705^ immunostaining in kidneys from 13 weeks old control, i*Pkd1*^ΔTub^ and i*Pkd1*^ΔTub^; i*Ccl2*^ΔTub^ mice. Scale bar: 50µm. **(B, C, E, F)** Each dot represents one individual mouse. One-way ANOVA followed by Tukey-Kramer test, ns: not significant, \**P*<0.05, \*\**P*<0.01, \*\*\**P*<0.001.

Collectively, these results indicate that, although primary cilia are direct positive regulators of STAT3 phosphorylation on Y705 *in vitro*, cilia dependent STAT3 activation in polycystic kidneys is mainly mediated through paracrine mechanisms requiring CCL2 dependent macrophage recruitment.

### Tubule specific *Stat3* inactivation does not prevent pre-cystic kidney growth in response to *Pkd1* disruption

How CCL2 dependent macrophage recruitment promotes cyst growth remains unclear in ADPKD. As STAT3 activation in tubular cells has been proposed to promote cyst growth, we investigated whether STAT3 activation in renal tubules is instrumental in ADPKD progression. To this aim, we crossed i*Pkd1*^Δtub^ with *Stat3*^flox/flox^ mice and derived mice allowing the inactivation of *Pkd1* alone (i*Pkd1*^Δtub^) or in association with *Stat3* (i*Pkd1*^Δtub^; i*Stat3*^Δtub^) specifically in tubular cells upon doxycycline treatment. At 13 weeks of age (7 weeks after the completion of doxycycline treatment), western blot analysis of kidney homogenates revealed a significant reduction of STAT3 protein in kidneys from i*Pkd1*^Δtub^; i*Stat3*^Δtub^ mice as compared to i*Pkd1*^Δtub^ mice (Figure 3A, B). STAT3 immunolabelling confirmed the loss of STAT3 protein in renal tubular cells from i*Pkd1*^Δtub^; i*Stat3*^Δtub^ mice (Figure 3C). In line with this observation, *Stat3* tubular inactivation in polycystin deficient mice markedly lessened *Socs3* induction (Figure 3D). At this time point, both i*Pkd1*^Δtub^ and i*Pkd1*^Δtub^; i*Stat3*^Δtub^ mice displayed kidney enlargement but overt polycystic kidneys was only observed in few i*Pkd1*^Δtub^ mice (Figure 3E-G). Although i*Pkd1*^Δtub^; i*Stat3*^Δtub^ kidneys tend to be smaller and less variable than i*Pkd1*^Δtub^ kidneys, this difference did not reach statistical significance. Collectively, these results demonstrate that *Stat3* is dispensable for the early kidney growth.

**Figure 3:**
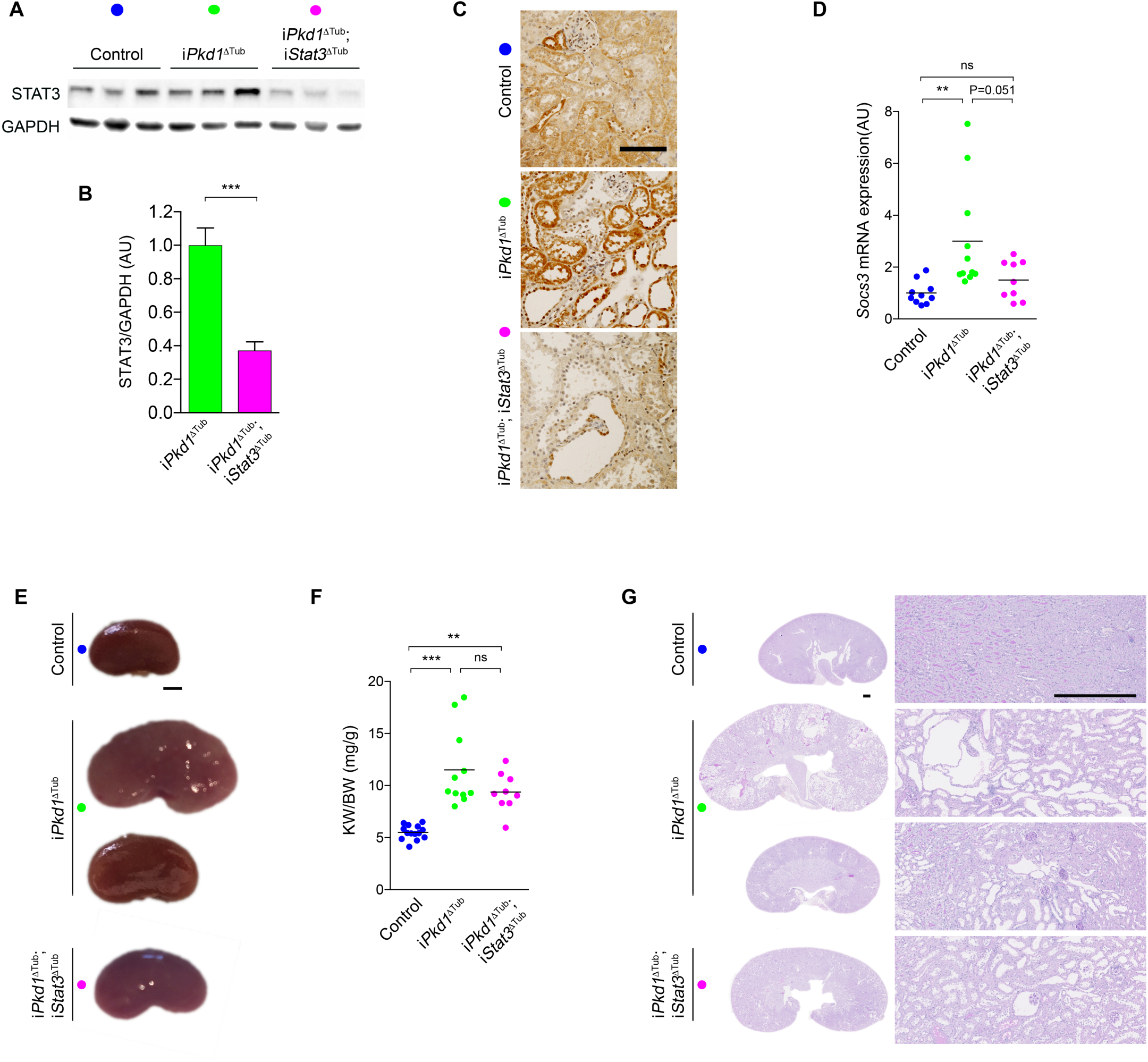
Tubule specific *Stat3* inactivation does not prevent early kidney growth in response to *Pkd1* disruption. **(A-B)** Representative western blot **(A)** and quantification **(B)** of STAT3 in kidney homogenates from 13 weeks old control, i*Pkd1*^ΔTub^ and i*Pkd1*^ΔTub^; i*Stat3*^ΔTub^ mice (7 weeks after the completion of doxycycline treatment). Each lane indicates one individual mouse. Bars are mean ± SEM of 6 mice per group. AU: arbitrary units. Two tailed *t* test, \*\*\**P*<0.0001. **(C)** STAT3 immunostaining of kidney sections from 13 weeks old control, i*Pkd1*^ΔTub^ and i*Pkd1*^ΔTub^; i*Stat3*^ΔTub^ mice. Representative images of 3 mice per group. Scale bar: 100µm. **(D)** Quantification of *Socs3* mRNA abundance in kidneys from 13 weeks old control, i*Pkd1*^ΔTub^ and i*Pkd1*^ΔTub^; i*Stat3*^ΔTub^ mice. **(E)** Representative kidneys from 13 weeks old control, i*Pkd1*^ΔTub^ and i*Pkd1*^ΔTub^; i*Stat3*^ΔTub^ mice. Scale bar: 2 mm. **(F)** Kidney weight to body weight ratio (KW/BW) at 13 weeks. **(G)** Representative Periodic Acid–Schiff (PAS) stained kidney sections from 13 weeks old control, i*Pkd1*^ΔTub^ and i*Pkd1*^ΔTub^; i*Stat3*^ΔTub^ mice at 13 weeks. Scale bar: 0.5mm. **(D, F)** Each dot represents one individual mouse. One-way ANOVA followed by Tukey-Kramer test, ns: not significant, \*\**P*<0.01, \*\*\**P*<0.001.

### Tubule specific *Stat3* inactivation does not prevent cystic kidney growth nor kidney function decline in response to *Pkd1* disruption

To assess the impact of loss of *Stat3* function on overt cystogenesis and kidney function decline, we let i*Pkd1*^Δtub^ and i*Pkd1*^Δtub^; i*Stat3*^Δtub^ mice age for 18 weeks (12 weeks after the completion of doxycycline treatment). Two over sixteen i*Pkd1*^Δtub^ and two over twenty i*Pkd1*^Δtub^; i*Stat3*^Δtub^ mice but none of the control mice died before the completion of the study (between 16 and 18 weeks; Figure 4A). i*Pkd1*^Δtub^ and i*Pkd1*^Δtub^; i*Stat3*^Δtub^ mice displayed a parallel modest decline in body weight in comparison to control mice at sacrifice (Figure 4B). Macroscopic inspection revealed more irregular kidneys in i*Pkd1*^Δtub^; i*Stat3*^Δtub^ than in i*Pkd1*^Δtub^ mice (Figure 4C), but overall, both i*Pkd1*^Δtub^ and i*Pkd1*^Δtub^; i*Stat3*^Δtub^ mice developed similar massive kidney enlargement as compared to control mice (Figure 4D). Microscopic inspection of the kidneys revealed that *Stat3* deletion resulted in a slight but significant decrease of cystic area (Figure 4E, F). In line with this modest effect on cyst burden, *Stat3* disruption not only reduced the expression of the STAT3 transcriptional targets *Socs3* and *Lcn2*, but also decreased the expression of *Havcr1* transcript, which encodes KIM-1, a marker of tubular injury that positively correlates with cyst burden in ADPKD patients^30^ (Figure 4G-I). In spite of this reduction in cyst burden both i*Pkd1*^Δtub^ and i*Pkd1*^Δtub^; i*Stat3*^Δtub^ mice displayed a similar increase in plasma creatinine and urea compared to control mice (Figure 4J, K), indicating that *Stat3* tubular deletion did not prevent polycystic kidney function decline. Of note, the more severe reductions in kidney function were observed in i*Pkd1*^Δtub^; i*Stat3*^Δtub^ animals (Figure 4J, K). These results demonstrate that tubular STAT3 plays only a marginal role in cyst growth in ADPKD and that *Stat3* tubular deletion does not prevent kidney function decline.

**Figure 4:**
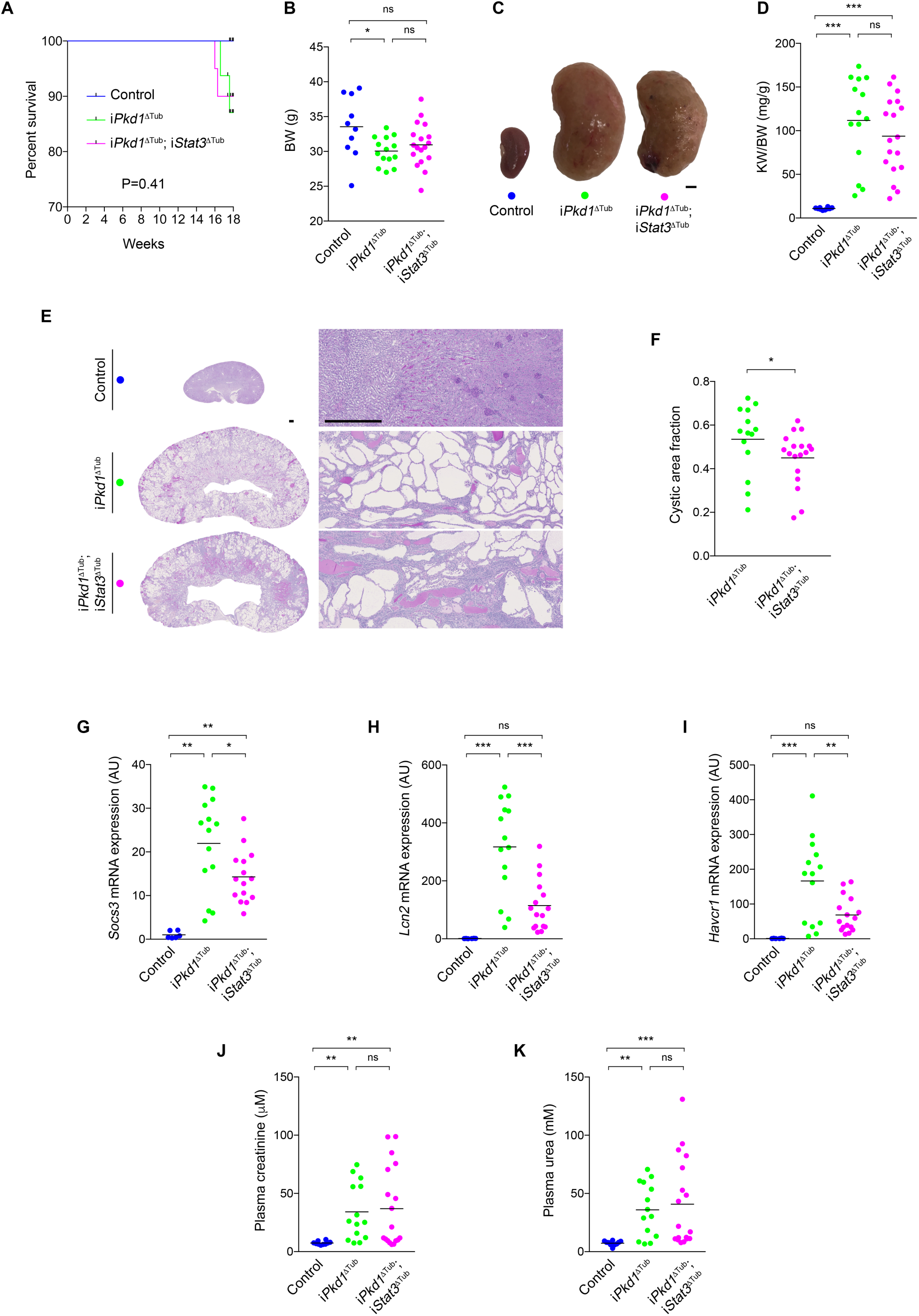
Tubule specific *Stat3* inactivation does not prevent cystic kidney growth nor kidney function decline in response to *Pkd1* disruption. **(A)** Kaplan Meyer survival curves of control, i*Pkd1*^ΔTub^ (n=16) and i*Pkd1*^ΔTub^; i*Stat3*^ΔTub^ (n=20) mice. **(B)** Body weight (BW) of 18 weeks old control, i*Pkd1*^ΔTub^ and i*Pkd1*^ΔTub^; i*Stat3*^ΔTub^ animals (12 weeks after the completion of doxycycline treatment). **(C)** Representative kidneys from the same animals. Scale bar: 2 mm. **(D)** Kidney weight to body weight ratio (KW/BW) at 18 weeks. **(E-F)** Representative PAS staining of kidneys sections **(E)** and measurement of cysts area fraction **(F)** from 18 weeks old control, i*Pkd1*^ΔTub^ and i*Pkd1*^ΔTub^; i*Stat3*^ΔTub^ mice. Scale bar: 0.5 mm. Mann-Whitney test, * *P*<0.05. **(G-I)** Quantification of *Socs3* **(G)**, *Lcn2* **(H)** and *Havcr1* **(I)** mRNA abundance in kidneys from 18 weeks old control, i*Pkd1*^ΔTub^ and i*Pkd1*^ΔTub^; i*Stat3*^ΔTub^ mice. **(J-K)** Measurement of plasma creatinine **(J)** and urea **(K)** from 18 weeks old control, i*Pkd1*^ΔTub^ and i*Pkd1*^ΔTub^; i*Stat3*^ΔTub^ animals. **(B, D, F, G-K)** Each dot represents one individual mouse. **(B, D, G-K)** One-way ANOVA followed by Tukey-Kramer test, ns: not significant, \**P*<0.05, \*\**P*<0.01, \*\*\**P*<0.001.

### Tubule specific *Stat3* inactivation increases polycystic kidney inflammation

The fact that i*Pkd1*^Δtub^; i*Stat3*^Δtub^ mice presented severe renal function impairment in spite of a reduction in cyst burden suggested that *Stat3* deletion could somehow worsen kidney damage without accelerating cyst growth. Indeed, we were surprised to observe a notable increase in interstitial inflammation in i*Pkd1*^Δtub^; i*Stat3*^Δtub^ mice (Figure 4G). Repeated semi-quantitative blinded assessment of the severity of the inflammatory infiltrates confirmed this difference (Figure 5A). Immunolabelling identified both T cells (CD3+) and macrophages (F4/80+) in the large inflammatory areas of i*Pkd1*^Δtub^; i*Stat3*^Δtub^ mice, while neutrophils (LY6C+) were rarely observed (Figure 5B). Consistently, we observed that *Stat3* inactivation was associated with a significant increase in the renal abundance of *Cd3* (expressed by T cells) and *Ccr2* (expressed by infiltrating macrophages) transcripts (Figure 5C, D).

**Figure 5:**
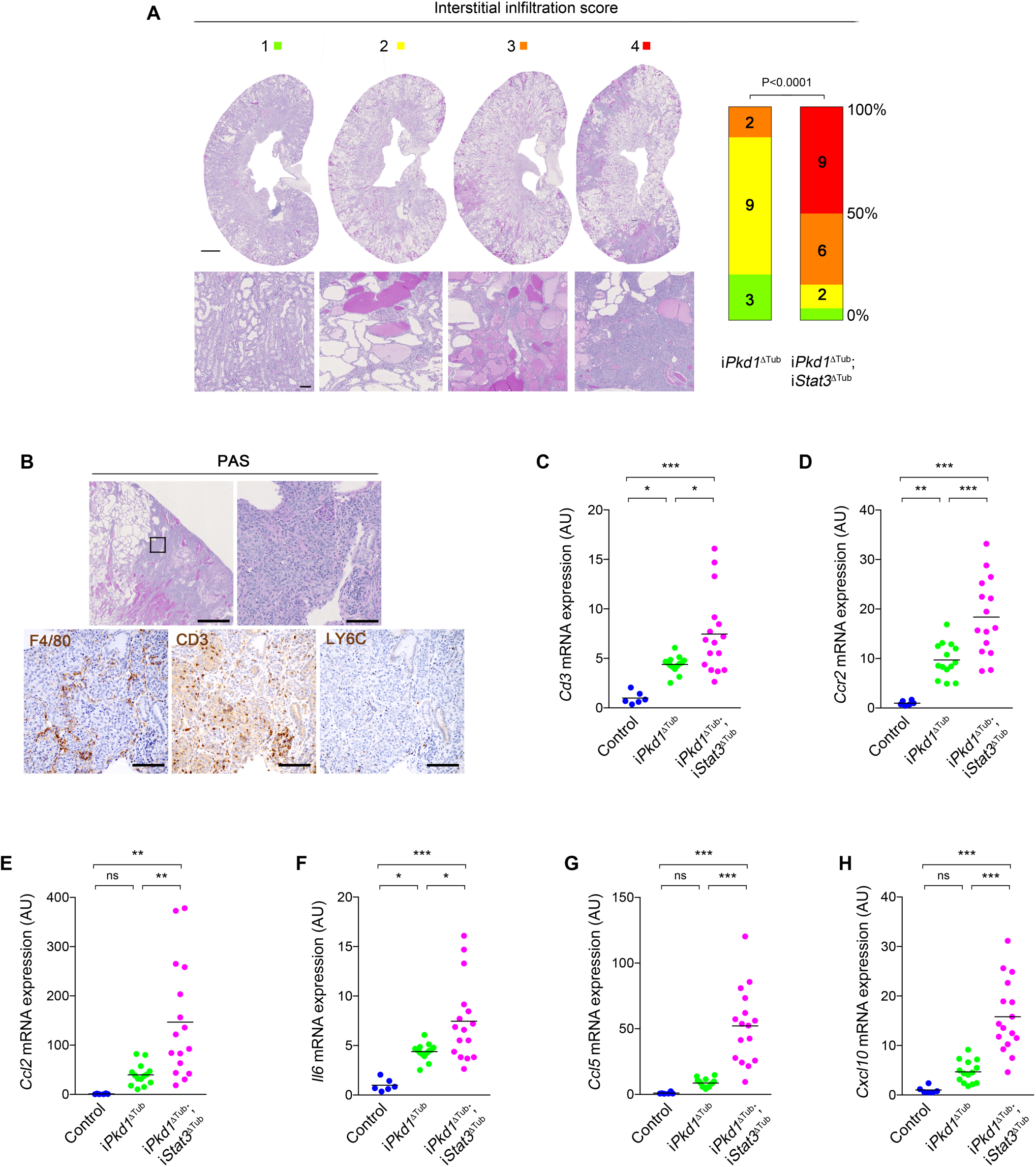
Tubule specific *Stat3* inactivation increases polycystic kidney inflammation. **(A)** Representative examples of interstitial inflammation scoring and distribution of renal interstitial inflammation scores in PAS stained kidneys from 18 weeks old i*Pkd1*^ΔTub^ and i*Pkd1*^ΔTub^; i*Stat3*^ΔTub^ mice. Numbers in the bars indicate number of mice per score; P is P value from Mann-Whitney test. **(B)** Representative images of PAS, F4/80, CD3 and LY6C immunostaining of kidney sections from 18 weeks old i*Pkd1*^ΔTub^; i*Stat3*^ΔTub^. Scale bars: 1mm for upper left panel and 0.1mm for other panels. **(C-H)** Quantification of *Cd3* **(C)**, *Ccr2* **(D)**, *Ccl2* **(E)**, *Il6* **(F)**, *Ccl5* **(G)** and *Cxcl10* **(H)** mRNA abundance in kidneys from 18 weeks old control, i*Pkd1*^ΔTub^ and i*Pkd1*^ΔTub^; i*Stat3*^ΔTub^ mice. Each dot represents one individual mouse. One-way ANOVA followed by Tukey-Kramer test, ns: not significant, \**P*<0.05, \*\**P*<0.01, \*\*\**P*<0.001.

A heightened inflammatory response has been observed after STAT3 ablation in hematopoietic cells, osteoblasts or keratinocytes^31,32^. In macrophages, the proinflammatory consequences of *Stat3* deletion are linked to an excessive activation of NF-κB signaling resulting in increased expression of NF-κB target genes^33^. We therefore investigated the impact of *Stat3* deletion on the expression of selected NF-κB targets. While *Pkd1* loss of function led to an increase in the abundance of *Ccl2*, *Il6*, *Ccl5 and Cxcl10* transcripts, *Stat3* inactivation amplified this phenomenon (Figure 5E-H). This was particularly evident for *Ccl5* and *Cxcl10*, whose expression was 6 and 3 fold higher in i*Pkd1*^Δtub^; i*Stat3*^Δtub^ than in i*Pkd1*^Δtub^ kidneys, respectively. In contrast, we did not detect any significant difference in *Tnfa*, *Il1b* or *Cxcl3* mRNA abundance between i*Pkd1*^Δtub^ and i*Pkd1*^Δtub^; i*Stat3*^Δtub^ kidneys (**Supplementary Figure 2**), indicating that *Stat3* deletion led to the upregulation of specific inflammatory mediators. In macrophages, STAT3 anti-inflammatory effects have been linked to the transcriptional repression of the E2 ubiquitin-conjugating enzyme UB2N, which stabilizes NF-κB signaling resulting in the nuclear accumulation of the transcription factor RELA^32^. However, contrary to its published effect in macrophages, tubule specific *Stat3* inactivation was not associated with an increase in *Ub2n* mRNA abundance in *Pkd1* deficient kidneys, suggesting alternative mechanisms (**Supplementary Figure 2**).

To investigate if STAT3 directly functions as a repressor of NF-κB induced proinflammatory cytokine in tubular cells, we analyzed the impact of STAT3 depletion on the induction of *Il6*, *Ccl2*, *Ccl5* and *Cxcl10* expression by LPS, a major activator of NF-κB, in tubular mIMCD-3 cells. While we were not able to detect *Il6* transcript in those cells, LPS treatment induced a robust induction of *Ccl2*, *Ccl5* and *Cxcl10* transcripts (Figure 6A-C). Strikingly, STAT3 depletion led to a proportional increase in the abundance of *Ccl5* and *Cxcl10* mRNA in response to LPS (Figure 6A, B). *Ccl2* mRNA levels showed the same trend as *Ccl5* and *Cxl10*, but statistically significant increase was restricted to the cell line with a profound STAT3 depletion (Figure 6C). Interestingly, LPS treatment led to STAT3 phosphorylation and nuclear accumulation, which was reduced by STAT3 depletion (Figure 6A, B). Contrary to macrophages, STAT3 depletion in mIMCD-3 did not increase the nuclear accumulation of RELA (Figure 6D, E). These results indicate that STAT3 dampens *Ccl5* and *Cxcl10* induction by LPS without affecting RELA nuclear accumulation. Collectively, these results indicate that STAT3 represses the expression of proinflammatory cytokines and restricts immune cell infiltration in ADPKD.

**Figure 6:**
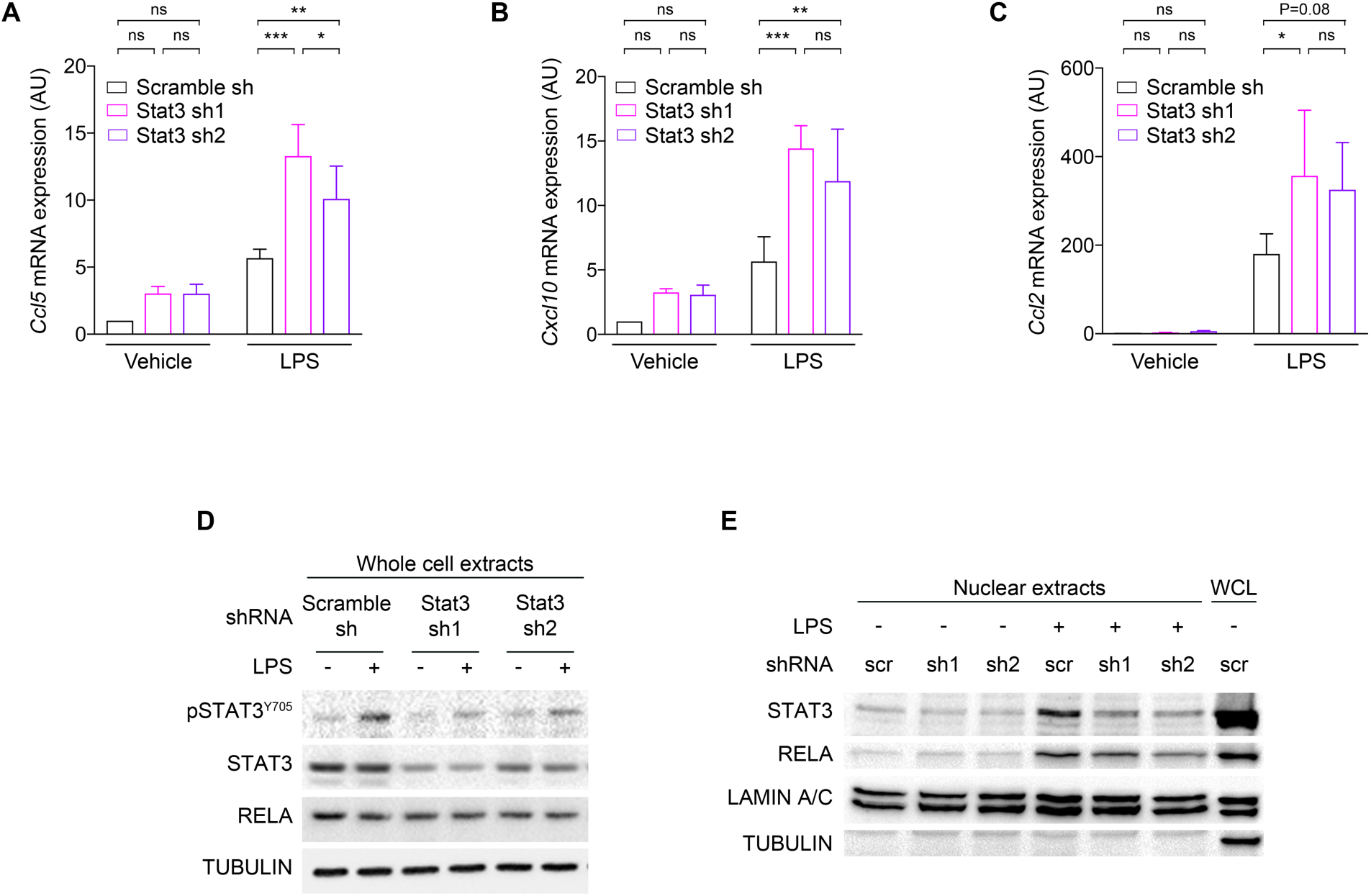
*Stat3* inactivation in tubular cells increases inflammatory cytokine transcriptional response. **(A-C)** Quantification of *Ccl2* **(A)**, *Ccl5* **(B)** and *Cxcl10* **(C)** mRNA abundance in mIMCD3 cells stably expressing shRNAs targeting STAT3 (sh1 and sh2) or a scramble control shRNA treated with lipopolysaccharide (LPS) or vehicle for 24 hours (24h). Bars are mean ± SEM of 8 independent experiments. Two-way paired ANOVA followed by Tukey-Kramer test, ns: not significant, \**P*<0.05, \*\**P*<0.01, \*\*\**P*<0.001. **(D)** Western blot of STAT3 phosphorylation and RELA expression in whole cell lysates from scramble shRNA (scr) and STAT3 shRNA (sh1 and 2) mIMCD3 cells, 24h after LPS or vehicle treatment. Representative western blot of 3 independent experiments. **(E)** Western blot of STAT3 and RELA expression in nuclear extracts from mIMCD3 cells expressing the indicated shRNAs, 24h after LPS or vehicle treatment. Whole cell lysates (WCL) served as a positive control for tubulin labelling. Representative western blot of 4 independent experiments.

## DISCUSSION

While multiple intrinsic defects in tubular cells homeostasis are observed in ADPKD^34–36^, accumulating evidence demonstrates that interstitial inflammation also plays an important role in the disease^6–8,10,37,38^. Yet, the nature of the factors mediating communication between tubular and immune cells and their impact on disease expression are not well-characterized. Two distinct pathways have been shown to promote macrophage accumulation in polycystic kidneys. First, polycystin deficiency triggers monocyte migration to the kidney through the induction of CCL2^7,10^. Second, kidney injury has been shown to promote cyst progression by triggering CSF1 dependent resident macrophage proliferation^37^. In turn, infiltrating macrophages drive tubular cell proliferation, possibly by releasing arginine metabolites^8^. In this context, our results unravel an unanticipated anti-inflammatory feedback loop orchestrated by STAT3 in ADPKD.

Previous reports have documented a robust activation of STAT3 in cyst lining epithelial cells from ADPKD patients and related murine models^19,20,39^. While *in vitro* results suggested that cleaved polycystin 1 may directly activate STAT3^39,40^, the mechanisms governing STAT3 activation in cystic tubular cells remained poorly understood. Our results indicate that the bulk of STAT3 activation in response to *Pkd1* disruption *in vivo* occurs through a non-cell autonomous process requiring a cilia dependent induction of CCL2. Acting on its receptor CCR2, CCL2 is a major driver of macrophage recruitment, which occurs at an early step of the disease^7,8,38^. Thus, our results imply that infiltrating macrophages directly or indirectly promote STAT3 activation in tubular cells. Such non-cell autonomous activation of STAT3 by neighboring macrophages have been documented in distinct settings including cancers^41–45^, heart failure^27^ or pancreatitis^46^. Different molecular pathways have been reported, including the direct expression of paracrine activator of STAT3 such as oncostatin M^27^, interleukin 6^47^, interleukin 8^42^ and TGFβ^46^ or the indirect up-regulation of STAT3 activating receptors^48^, mirroring the multiplicity of STAT3 activation mechanisms.

In most extra-renal settings, paracrine activation of STAT3 by macrophages plays a detrimental effects in the disease, promoting tissue injury^27^ or tumor growth/metastasis^41–45^. In contrast to these situations, our data indicate that tubular *Stat3* is mostly dispensable for cyst growth in ADPKD, but dampens immune cell infiltration of polycystic kidneys. The absence of a notable effect of *Stat3* tubular deletion on kidney enlargement and kidney function decline in our model sharply contrasts with the positive impact of STAT3 inhibitory compounds previously reported in *Pkd1* mutant mice^19,20^. This discrepancy may be caused by off-target effects or the consequence of inadequate target specificity of these drugs^20,21,49^. Alternatively, the beneficial effect of these drugs may be linked to *Stat3* inhibition in non-tubular cells. Indeed, pericystic interstitial cells also display STAT3 activation in ADPKD. Importantly, different interstitial cell types have been shown to affect cyst growth, including macrophages^6,8^ and CD8+ T cells^38^. Interestingly, STAT3 activation has been reported in pericystic macrophages in ADPKD and STAT3 inhibitory compounds reduce the ability of cultured macrophages to induce tubular cell proliferation through paracrine signaling *in vitro*^50^. It is therefore plausible that the therapeutic effects of STAT3 inhibitory drugs are linked to the inhibition of STAT3 in macrophages. The proper evaluation of STAT3 functions in macrophage in ADPKD represents however a complicated task: conditional *Stat3* inactivation in macrophages leads to the spontaneous development of inflammatory bowel disease precluding the use of such approach in a murine model of ADPKD^51^. Thus, our results do not rule out that STAT3 inhibition may have a benefit in ADPKD patients through its impact on non-tubular cells.

Unexpectedly, we observed that *Stat3* deletion in renal tubules markedly increased immune cell infiltration of polycystic kidneys. Mechanistically, STAT3 depletion resulted in the up-regulation of *Ccl5* and *Cxcl10* transcripts in polycystic kidneys and in cultured tubular cells. While surprising, this observation is reminiscent of the well characterized anti-inflammatory functions of STAT3 in macrophages. In those cells, STAT3 activation in response to IL10 restricts NF-κB dependent proinflammatory signaling^31,32^. The molecular bases of this anti-inflammatory effect involves the transcriptional regulation of factor(s) that modulate(s) NF-κB signaling^31^. More specifically, the transcriptional repression of UB2N ubiquitin ligase by STAT3 appears to play a major role in restricting NF-κB signaling in macrophages^32^. Indeed, in macrophages, *Stat3* inactivation increases UB2N abundance, which in turn stabilizes NF-κB signaling and promotes RELA nuclear accumulation. Of note, STAT3 anti-inflammatory functions have also been documented in non-immune cells. Indeed, the deletion of STAT3 in intestinal epithelial cells leads to an exacerbation inflammatory response after intestinal injury^17^. We did not fully investigate the underlying mechanisms of STAT3 anti-inflammatory function in tubular cells, but our results indicate some specificities: contrary to STAT3 anti-inflammatory functions in macrophages, the anti-inflammatory effect of STAT3 in tubular cells is not mediated by the repression of *Ub2n* transcript, nor is it associated with RELA nuclear accumulation. While the molecular intermediates allowing STAT3 to restrict polycystic kidney inflammation remain unclear, our results clearly expand STAT3 anti-inflammatory function to tubular cells. They further suggest that this apparently conserved function of STAT3 is related to distinct tissue specific mechanisms.

A striking aspect of our study is that tubular *Stat3* disruption decoupled renal inflammation from cyst growth. Indeed, previous studies have consistently shown that reducing renal inflammation through macrophage depletion or CCL2/CCR2 inhibition led to a proportional decrease in cyst burden. Why increased macrophage infiltration does not sustain cyst growth in *Stat3* deficient animals remain unclear. It may indicate a direct effect of STAT3 on tubular cell proliferation as suggested by some *in vitro* studies^19,52^. Considering the role played by CD8+ T cells in restricting cyst growth^38^, the massive increased in T cells infiltration induced by *Stat3* inactivation could also be involved.

The identification of V2R inhibition as a valid target to reduce cyst growth in human has opened the area of specific therapy in ADPKD^53,54^. While this represents an important cornerstone on the road to cure the disease, V2R inhibition has only a modest efficacy and important side effects. Thus, the search for more potent and better tolerated drugs remains central to the field^55^. Over the last 20 years, joint research efforts allowed the identification of a large panel of pathways that are activated in growing cysts, thereby providing multiple potential target for molecular targeted therapies in ADPKD^56^. While concordant evidence identified STAT3 as an interesting candidate, robust evaluation of STAT3 activation mechanisms and functions in ADPKD was lacking. In this context, our study, which relay on state of the art model of tubular specific inactivation of *Stat3* and *Pkd1* in adult mice, clearly indicates that tubular STAT3 is not a critical driver of cyst formation. More generally, our results exemplify the critical contribution of complex genetic conditional models to the rigorous evaluations of potential therapeutic targets in ADPKD.

## ACKNOWLEDGMENTS

We thank the animal facility (LEAT, SFR Necker, INSERM US24, Paris, France), the histology facility (SFR Necker, INSERM US24, Paris, France) and the mice physiology facility (Cordelier research center, Paris, France) for technical assistance.

Amandine Viau was supported by FRM ARF20150934110, E. Wolfgang Kuehn was supported by DFG 1504/5-1, Fabiola Terzi & Frank Bienaimé were supported by Institut National de la Santé et de la Recherche Médicale, Université Paris Descartes, Assistance Publique Hôpitaux de Paris and Agence Nationale de la Recherche.

## AUTHOR CONTRIBUTIONS

A.V and F.B designed the study; A.V, M.B, A.A, C.N and F.B performed experiments and analyzed data; A.V, E.W.K, F.T and F.B drafted and revised the manuscript; all authors approved the final version of the manuscript.

---

**Supplementary Figure 1: Primary cilia promote STAT3 phosphorylation in tubular cells *in vitro*.**

**(A, B)** Representative western blot **(A)** and quantification **(B)** of STAT3 phosphorylation in whole cell lysates from MDCK cells expressing inducible shRNA against *Kif3a* (Kif3a-i), *Ift88* (Ift88-i) or the non-relevant control luciferase (Luci-i) after 10 days of tetracyclin treatment (+ Tet). Bars are mean ± SEM of 4 independent experiments. Paired *t* test, ns: not significant, \**P*<0.05.

**Supplementary Figure 2: STAT3 disruption has a selective impact on inflammatory cytokine expression in ADPKD.**

**(A-D)** Quantification of *Tnfa* **(A)**, *Il1b* **(B)**, *Cx3cl1* **(C)**, and *Ub2n* **(D)** mRNA abundance in kidneys from control, i*Pkd1*^ΔTub^ and i*Pkd1*^ΔTub^; i*Stat3*^ΔTub^ mice at 18 weeks. Each dot represents one individual mouse. One-way ANOVA followed by Tukey-Kramer test, ns: not significant, \*\**P*<0.01, \*\*\**P*<0.001.

