## Supplementary material for "Tubular STAT3 limits renal inflammation in autosomal dominant polycystic kidney disease"

### Table of contents

Supplementary Figure 1: Primary cilia promote STAT3 phosphorylation in tubular cells *in vitro*.

Supplementary Figure 2: STAT3 disruption has a selective impact on inflammatory cytokine expression in ADPKD.

Supplementary Table 1: Primers used for qRT-PCR.

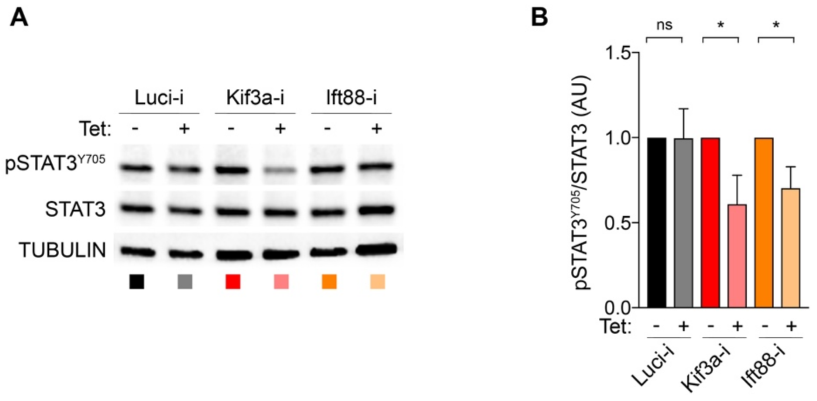


**Supplementary Figure 1: Primary cilia promote STAT3 phosphorylation in tubular cells *in vitro*.**

**(A, B)** Representative western blot **(A)** and quantification **(B)** of STAT3 phosphorylation in whole cell lysates from MDCK cells expressing inducible shRNA against *Kif3a* (Kif3a-i), *Ift88* (Ift88-i) or the non-relevant control luciferase (Luci-i) after 10 days of tetracyclin treatment (+ Tet). Bars are mean ± SEM of 4 independent experiments. Paired *t* test, ns: not significant, **P*<0.05.

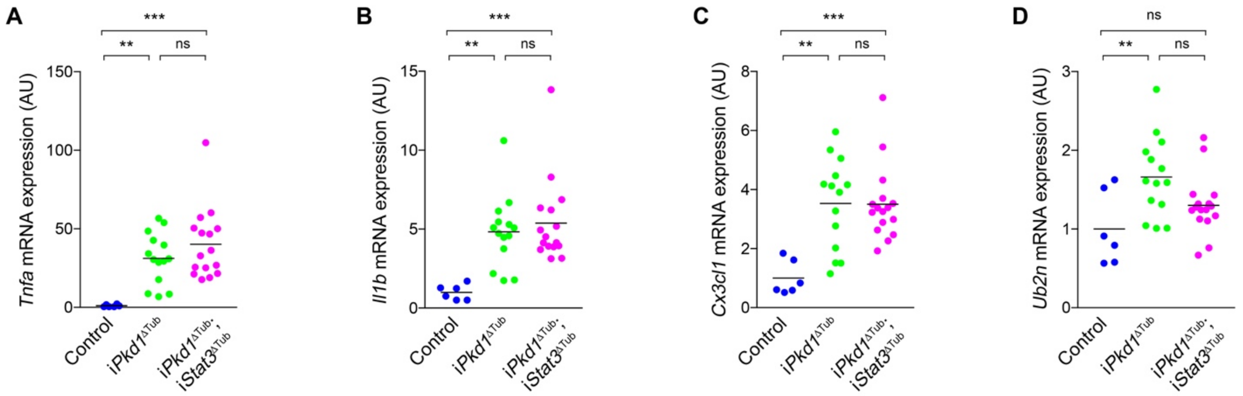


**Supplementary Figure 2: STAT3 disruption has a selective impact on inflammatory cytokine expression in ADPKD.**

**(A-D)** Quantification of *Tnfa* **(A)**, *Il1b* **(B)**, *Cx3cl1* **(C)**, and *Ub2n* **(D)** mRNA abundance in kidneys from control, i*Pkd1*$^{\Delta Tub}$ and i*Pkd1*$^{\Delta Tub}$; i*Stat3*$^{\Delta Tub}$ mice at 18 weeks. Each dot represents one individual mouse. One-way ANOVA followed by Tukey-Kramer test, ns: not significant, ***P*<0.01, ****P*<0.001.

**Supplementary table 1: primers used for RT-qPCR**

| | Forward Primer (5' to 3') | Reverse Primer (5' to 3') |
|---|---|---|
| Primers for *Canis Lupus* | | |
| *Ccl2* | CTGCTGCTATACACTCACCAATA | TCAGCACAGATCTCCTTGTTTAG |
| *Gapdh* | CATGTTTGTGATGGGCGTGAACCA | TTTGGCTAGAGGAGCCAAGCAGTT |
| Primers for *Mus Musculus* | | |
| *Ccl2* | AGTAGGCTGGAGAGCTACAA | GTATGTCTGGACCCATTCCTTC |
| *Ccl5* | GCCCACGTCAAGGAGTATTT | CTTGAACCCACTTCTTCTCTGG |
| *Ccr2* | GCTCTACATTCACTCCTTCCAC | ACCACTGTCTTTGAGGCTTG |
| *Cd3e* | AAGCCTGTGACCCGAGGAA | TGCGGATGGGCTCATAGTCT |
| *Cx3cl1* | GGAAAGAAACGTGGTCCAGA | GGAAAGAAACGTGGTCCAGA |
| *Cxcl10* | GGATGGCTGTCCTAGCTCTG | TGAGCTAGGGAGGACAAGGA |
| *Gapdh* | TGCACCACCAACTGCTTAG | TGGATGCAGGGATGATGTT |
| *Havcr1* | GAGAGTGACAGTGGTCTGTATTG | CCTTGTAGTTGTGGGTCTTCTT |
| *Hprt* | GGCCAGACTTTGTTGGATTTG | CGCTCATCTTAGGCTTTGTATTTG |
| *Il1* | GAGGACATGAGCACCTTCTTT | GCCTGTAGTGCAGTTGTCTAA |
| *Il6* | CTCTGGGAAATCGTGGAAATG | AAGTGCATCATGGTTGTTCAT |
| *Lcn2* | GGACCAGGGCTGTCGCTACT | GGTGGCCACTTGCACATTGT |
| *Ppia* | GGCTATAAGGGTTCCTCCTTTC | TTTCTCTCCGTAGATGGACCT |
| *Rpl13* | CTCATCCTGTTCCCCAGGAA | GGGTGGCCAGCTTAAGTTCTT |
| *Socs3* | CCACCCTCCAGCATCTTTGT | CAGGCAGCTGGGTCACTTTC |
| *Tnfa* | ATTCGAGTGACAAGCCTGTAG | TGAAGAGAACCTGGGAGT |
| *Ub2n* | CAGAACCAGTTCCTGGCATT | CAGTGCTGGGGACCACTTAT |
